# Automatic Three-Dimensional Reconstruction of Fascicles in Peripheral Nerves from Histological Images

**DOI:** 10.1101/2020.01.22.913251

**Authors:** Daniel Tovbis, Anne Agur, Jeremy P.M. Mogk, José Zariffa

## Abstract

Computational studies can be used to support the development of peripheral nerve interfaces, but currently use simplified models of nerve anatomy, which may impact the applicability of simulation results. To better quantify and model neural anatomy across the population, we have developed an algorithm to automatically reconstruct accurate peripheral nerve models from histological cross-sections. We acquired serial median nerve cross-sections from human cadaveric samples, staining one set with hematoxylin and eosin (H&E) and the other using immunohistochemistry (IHC) with anti-neurofilament antibody. We developed a four-step processing pipeline involving registration, fascicle detection, segmentation, and reconstruction. We compared the output of each step to manual ground truths, and additionally compared the final models to commonly used extrusions, via intersection-over-union (IOU). Fascicle detection and segmentation required the use of a neural network and active contours in H&E-stained images, but only simple image processing methods for IHC-stained images. Reconstruction achieved an IOU of 0.42±0.07 for H&E and 0.37±0.16 for IHC images, with errors partially attributable to global misalignment at the registration step, rather than poor reconstruction. This work provides a quantitative baseline for fully automatic construction of peripheral nerve models. Our models provided fascicular shape and branching information that would be lost via extrusion.

## 1. INTRODUCTION

Neural interfaces (NIs) are systems that serve to exchange information between target neural structures and artificial devices. NIs are used in neuroprosthetic systems aiming to restore sensorimotor function after damage to the nervous system, as well as in neuromodulation systems aiming to treat diseases through the alteration of regulatory neural signals. Applications of NIs implanted in the peripheral nervous system include: restoring movement after paralysis (1); creating prosthetic limbs with intuitive control and sensory feedback(2); and treating conditions such as bladder dysfunction(3), epilepsy(4), hypertension(5), as well as inflammatory and autoimmune disorders(6). Despite their potential benefits, widespread implementation of NIs in the peripheral nervous system still faces several obstacles, including damage to neural tissue, a lack of long-term stability, and low signal resolution(7). These issues may be solved, or at least mitigated, by improving the design of new NIs. Improvements may include use of new materials or electrode designs, which have the potential to increase the effectiveness and reliability of NIs.

An important part of the design process for new NI designs is computational modeling (8–10). To be useful, a model should contain sufficient detail to capture the relevant features of the system of interest. Unfortunately, many existing peripheral nerve models used to design and evaluate NIs have been based on simplified anatomy – either by extruding a single “realistic” cross-section, or by assuming fascicles possess circular and/or elliptical cross-sections (11–13). Recent studies have shown that differences in peripheral neural anatomy, such as fascicular branching, can significantly alter the characteristics of neural recordings and the conclusions drawn from a computational model. Using an anatomically accurate fascicular model can for example alter the relative amplitudes across electrode recording sites, affecting conclusions about selectivity (14). Complementing this finding, implanting an electrode before or after a fascicular branch has been shown to alter recording selectivity *in vivo* (15). Thus, computational models that accurately reflect fascicular anatomy will improve the validity and applicability of the conclusions, and may ultimately lead to improved NI designs(13).

An anatomically accurate model of a peripheral nerve can be constructed using data acquired from a variety of imaging modalities, including histological cross-sections, Micro-Computed Tomography (MicroCT), Optical Projection Tomography (OPT), or Magnetic Resonance Imaging (MRI) (14,16,17). To date, either fully manual or semi-automatic procedures have been used to reconstruct peripheral nerve models from image data (17–19). However, the internal anatomy of peripheral nerves varies across the population, so simply acquiring one fascicular model fails to capture the population-level variability that may be useful to inform NI design or surgical decision making (20). Considering the benefits of capturing population-level anatomical data, coupled with the rapidly increasing interest in peripheral NIs for multiple applications, this manual or semi-automatic construction of computational models becomes non-feasible.

This paper introduces an image processing pipeline that, given a set of histological cross-sections of a nerve, aims to automatically identify and segment fascicles, and reconstruct a 3D model of a nerve’s internal fascicular anatomy. This tool is intended to greatly facilitate the generation of peripheral nerve models, while improving their anatomic fidelity. These reconstructions can replace simplified extrusion models to produce more accurate simulation outputs, incorporate statistical information about the population, and thus better inform the design of future NIs. The system we developed ultimately proves to be a baseline for future development, supported by quantitative data.

## 2. MATERIALS AND METHODS

A four-step process was used to process images: 1) registration of consecutive slices; 2) fascicle detection; 3) fascicle segmentation; and 4) 3-dimensional reconstruction. Details of the data acquisition and processing steps follow below.

### 2.1. Sample Acquisition

Acquisition of high quality nerve samples will allow for generation of the best possible quality images for automatic registration and segmentation. Five median nerve specimens obtained from embalmed human cadaveric forearms were used. Exclusion criteria included any visible evidence of deformities, previous surgery, or pathology. Ethics approval was obtained from the University of Toronto, Health Sciences Research Ethics Board (#27210). The median nerve was chosen as it is both relatively easy to extract and relevant for the purpose of upper limb NI applications.

To obtain the nerve segments for histological analysis, all superficial tissues were removed to expose the flexor digitorum superficialis muscle (FDS) and median nerve from the medial epicondyle to the nerve’s entry point into the muscle belly. Next, the median nerve and FDS were excised proximally at the elbow and distally at the wrist. The excised specimen was placed in a tray and the median nerve dissected to expose the intramuscular nerve branches. (Figure 1, top)

**Fig. 1:**
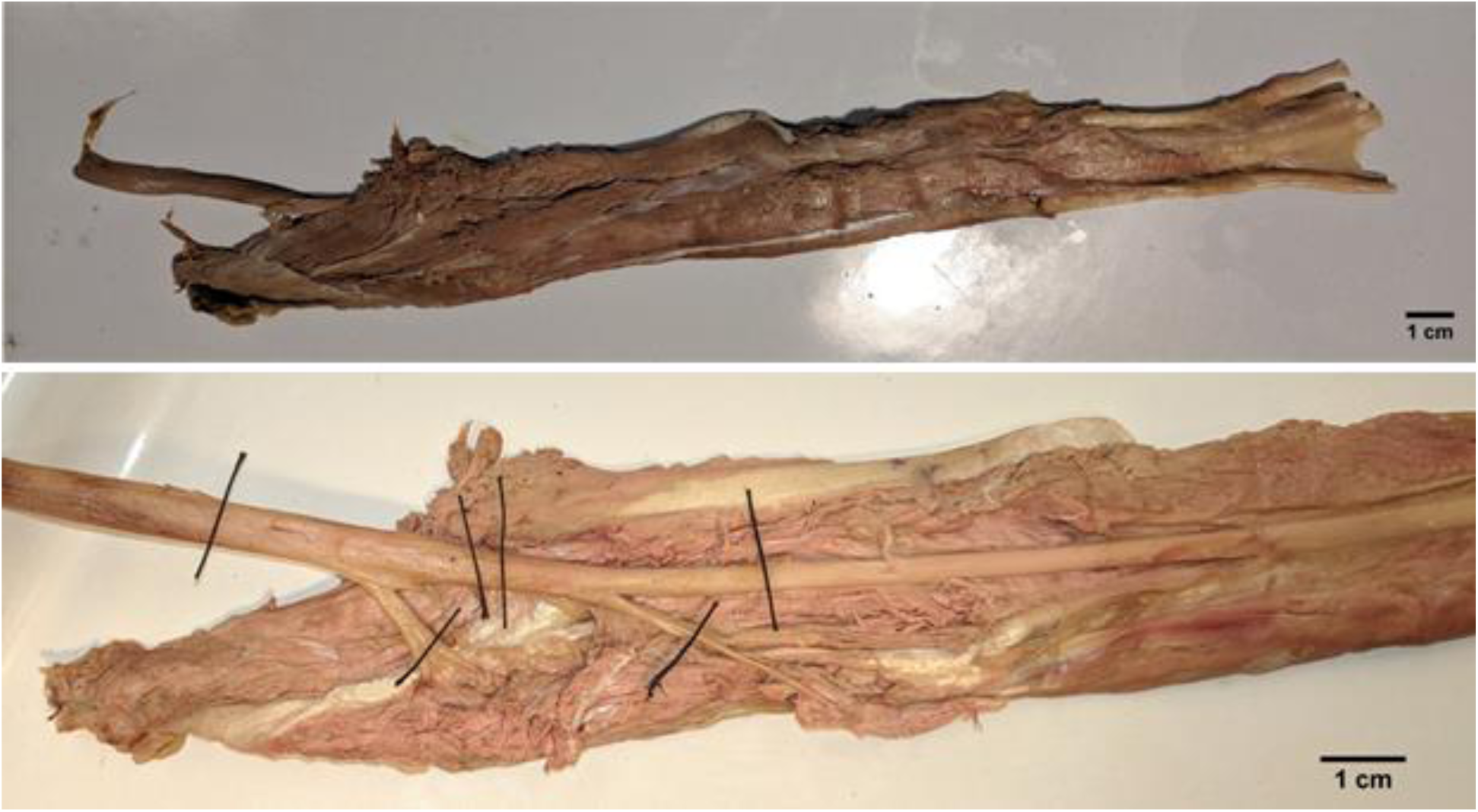
Top: A length of median nerve surrounded by flexor digitorum superficialis muscle excised from a cadaveric specimen. Bottom: The same sample, showing the nerve with the sections marked for extraction

For histological analysis, two median nerve segments were obtained from each of the five specimens. The proximal segment was obtained by ligating the nerve approximately 2 cm proximal and distal to the first bifurcation point (4 cm segments). The distal segment consisted of the same length of nerve proximal and distal to the second bifurcation point (Figure 1, bottom). The proximal and distal nerve segments from two specimens (n=4) were used to determine optimal staining methods, while the remaining nerve segments (n=6) were used for data collection.

### 2.2. Histology

All histological processing was performed by the Centre for Phenogenomics (Toronto, Ontario). The nerve segments were fixed in formalin, embedded in paraffin, and sectioned at 5μm thickness separated by 250μm intervals, resulting in 2-5 contiguous blocks of nerve slices per segment. From each specimen, either the proximal or the distal segment was stained with hematoxylin and eosin (H&E) and the other segment with anti-neurofilament antibody immunohistochemistry (IHC) (Figure 2). The sections were mounted on glass slides, with 3-5 slices per slide. Images were captured with an optical microscope (Olympus VS-120, Olympus Corporation, Tokyo, Japan) at 40x magnification. To facilitate further image processing, each slice was saved separately at 1.4x magnification (∼5183 pixels per inch) and stored as a lossless TIFF image. We collected a total of 130 images (10 blocks, 13 images per block) from the first specimen (65 H&E images from the proximal segment, 65 IHC images from the distal segment). A total of 144 images (12 blocks, 12 images per block) were obtained from the second (36 H&E, proximal segment/36 IHC, distal segment) and third (36 IHC, proximal segment/36 H&E, distal segment) specimens. The images obtained from one of the three segments stained with IHC showed no visible fascicles and was replaced. Any images that exhibited artifacts due to tissue processing were excluded (n=20). The final count of images used for testing the pipeline was 254 (Table 1).

**Fig 2:**
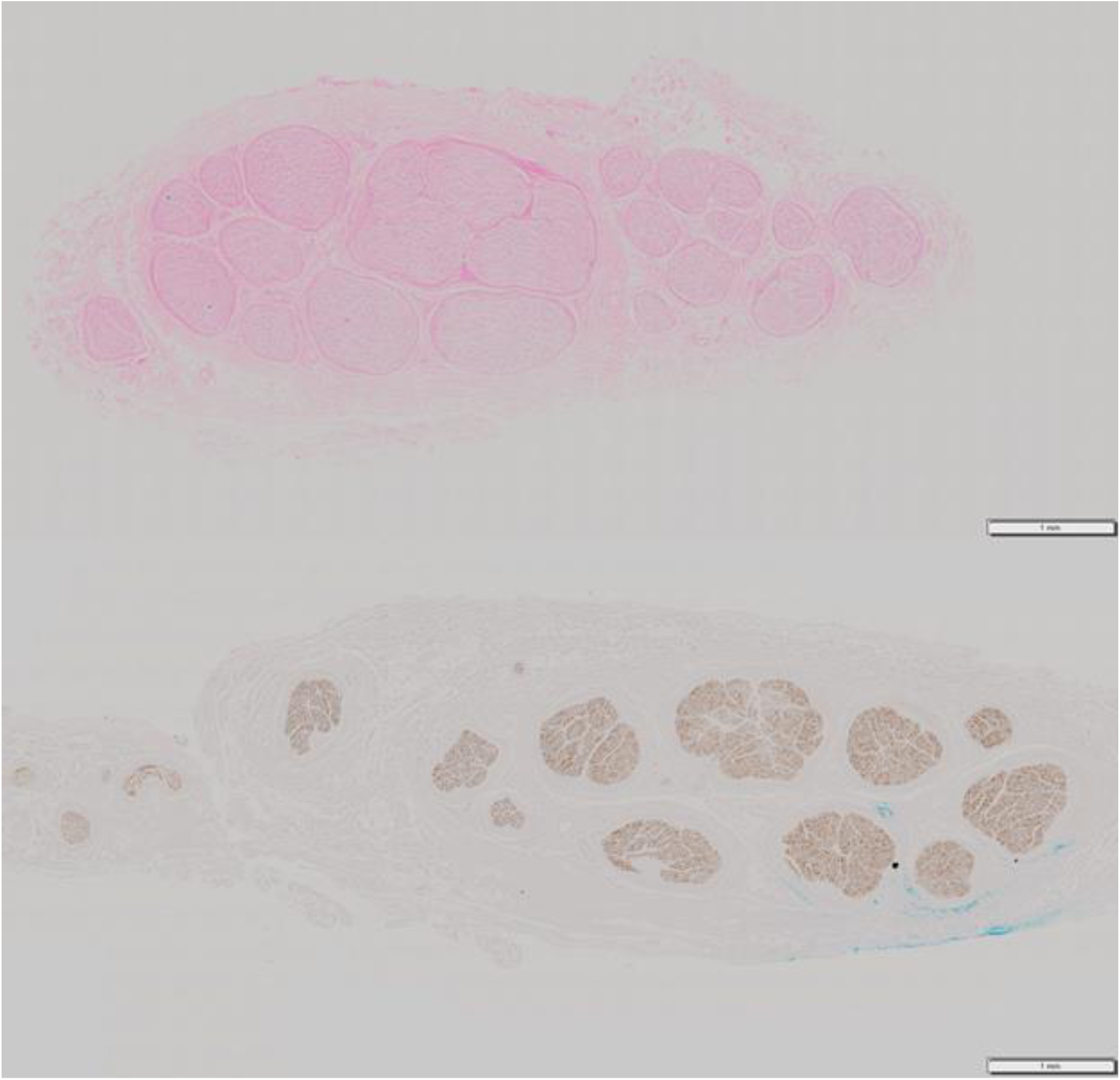
Two sample nerve slices, one stained with H&E (top) and the other with IHC (bottom).

**TABLE 1.** Summary of details related to nerve segments used to test the pipeline.

| Specimen | Segment | Stain | Contiguous Blocks | Slices per Block |
| --- | --- | --- | --- | --- |
| 1 | Proximal | HE | 5 | 9/13/13/13/12 |
| 1 | Distal | IHC | 5 | 9/11/11/12/10 |
| 2 | Proximal | HE | 3 | 12/12/12 |
| 2 | Distal | IHC | 3 | 12/12/12 |
| 3 | Distal | HE | 3 | 12/12/12 |
| 3 | Proximal | IHC | 3 | N/A (no neural tissue) |
| 4 | Distal | IHC | 2 | 16/17 |

### 2.3. Registration

Registration and all further processing was performed in MATLAB (r2018a, The MathWorks, Natick MA). Prior to registration, preprocessing steps for H&E stained images included an increase in local contrast (Edge Threshold 0.3, Enhancement 0.6) and a sharpening to emphasize fascicles and fascicle boundaries. The image was then converted to grayscale and processed via anisotropic diffusion filtering (5 iterations, exponential conduction method), a method which regularizes shape interiors while preserving edges (21). Morphological opening- and closing-by-reconstruction followed. Structural element sizes or erosion and dilation were set to disks of a radius of 7 pixels, according to the fascicle sizes in the images. Preprocessing for IHC images was minimal and involved only a conversion to greyscale.

Preprocessed images were aligned automatically using intensity-based image registration between consecutive slices with the stochastic gradient descent method (SGDM). We used a four-step registration method similar to the MIAQuant method (22), sequentially applying the registration four times. Each step used a different combination of registration types and image scale. These were 1) translation only, using images resized to 25% of their initial resolution 2) translation and rotation, using images resized to 50%, 3) translation, rotation, and scale, using images resized to 75% (H&E) or full resolution images (IHC) and 4) translation, rotation, scale, and shear using full resolution images. This multi-resolution procedure permits registration of gross image features first (minor details are lost when the image is scaled down), followed by progressive fine-tuning. Rotation produced black regions around the edges of the image. These were filled in with the background colour to prevent interference in registration.

### 2.4. Fascicle Detection

To identify fascicles stained with H&E, a pre-trained convolutional neural network using the VGG-16 network architecture was converted into a region-based convolutional neural network (RCNN) for fascicle detection and retrained using the MATLAB neural network toolbox on the Neuroscience Gateway (23). Detection created a set of bounding boxes roughly delineating the outer boundaries of the fascicles. Using a cross-validation process, the network was trained, in turn, using two of the three sets of nerve images and tested using the third. The three sets contained 1296, 2158, and 2388 annotated nerve fascicles. The three sets of images were manually annotated with fascicle bounding boxes, which served as the ground truth to evaluate the performance of the network. Two types of networks were tested; one using registered images for training, and the other using unregistered images. The parameters used for training the network were as follows: a batch size of 128, an initial learning rate of 1*10^-3^, a learning rate drop factor of 0.1, a learning rate drop period of 5 epochs, and a total training period of 10 epochs. Inputs to the RCNN were raw, unprocessed images. Since the IHC images showed high contrast between fascicular and non-fascicular tissue, a separate detection step was not necessary.

### 2.5. Fascicle Segmentation

For H&E slices, the bounding boxes proposed by the RCNN were used to generate ovals around each detected fascicle. The image outside of these ovals was removed to reduce the chance of mis-segmentation. A mask, slightly larger than the ovals generated by the RCNN, was then created for segmentation via Chan-Vese active contours(24). The Chan-Vese method used a smooth factor of 0.6 and a contraction bias of 0.5. The images were then pre-processed once again, using the same method as in the registration step. The contour then shrunk inwards until the border of the fascicle was detected, or 500 iterations passed. For the IHC slices, an automatic threshold generated by Otsu’s method (25), followed by image closing and hole-filling to account for non-uniform staining, was sufficient to produce a binary mask.

### 2.6. Reconstruction

The final step of generating a 3D model of the fascicular anatomy was accomplished by linearly interpolating between points on successive images within each block using a difference of distance maps (26). The algorithm connected shapes at different points on the z-axis by interpolating the distance between any given pixel and the object boundary along the z-axis. Shapes could be connected along the z-axis provided some overlap existed in the x-y plane. Two additional steps complemented the interpolation. The first step compensated for missed detections and mis-segmentations. Before reconstruction, the process checked the pixel indices of each fascicle on each image against those same indices on the next image. If no fascicle was detected at a particular index, but appeared in any future layer (indicating a missed detection or mis-segmentation), a fascicle was inserted at each missing index to prevent discontinuous fascicles (“hole fixing”, HF, compared to “no fixing”, NF). The second step was implemented to split erroneously merged fascicles and consisted of a watershed operation initialized using a distance map derived from the fascicle segmentation. An erosion and another watershed followed, intended to catch any large merged fascicles that the first watershed might have missed, though usually unnecessary. The fascicles were then dilated back to their original sizes This process produced one model for each block of nerve images, for a total of 21 fully automatic models.

### 2.7. Evaluation and Analysis

#### 2.7.1 Registration

Registration quality was calculated using two metrics: mean square error (MSE) and structural similarity (SSIM). MSE decreases and SSIM increases upon a successful registration. Net MSE and SSIM values, from before and after registration, were compared using Friedman’s test with a significance value of 0.05. To determine whether or not values were normally distributed, the Anderson-Darling test was used to indicate non-parametric values.

#### 2.7.2. Fascicle Detection

Fascicle detection in the H&E slices was tested by comparing automatically detected fascicle bounding boxes to the manually-labeled ground truth, using the F1-score (harmonic mean of precision and recall) to quantify the accuracy of the detector. An F1 score closer to 1 indicates high precision and recall. Any detected bounding box that overlapped a ground truth bounding box with an intersection-over-union (IOU) threshold of 50% was determined to be a true positive detection. To provide a point of comparison for this approach, the neural network detections were compared against Atherton-Kirby’s phase-coded method for circle detection(27). This method uses a modified version of the Circle Hough Transform to detect circles in images.

#### 2.7.3. Fascicle Segmentation

For both H&E and IHC slices, segmentation quality was tested by comparing the results of automatic segmentation to a manually segmented ground truth via IOU. Ground truth segmentations were acquired by manually labelling fascicles using MATLAB. When uncertainty existed as to whether or not a certain feature was indeed neural tissue, the original microscope slides were consulted (viewed at up to 40x magnification) to ensure that the stained feature was indeed a fascicle. Segmentation by active contours for the H&E slices was also compared to automatic thresholding using Otsu’s method and K-means clustering.

#### 2.7.4. Reconstruction

No objective ground truth regarding the 3D shape of the fascicles was available to this project, since 3D imaging (e.g., MicroCT, OPT) was not performed prior to histological sectioning. Thus, images were manually registered and combined with their manual segmentations to make fully manual models, which represented our “gold standard”. Our evaluation for the reconstruction involved calculating 3D IOU between the fully manual models and the following, for two random blocks from each segment (total of six H&E and six IHC blocks):

i. The corresponding fully automatic models.
ii. The manually registered images made for the fully manual models were automatically segmented (MRAS). This helped determine how differences in segmentation affected the reconstruction method.
iii. Automatically registered images were combined with manual segmentations (ARMS) to make a second set of semi-automatic models. This helped determine differences in registration affected the reconstruction method.

Along with the output of the automatic and semi-automatic pipelines, an extrusion (EX) generated from a manual segmentation of the first image in the block was compared to the fully manual models.

## 3. RESULTS

We tested the pipeline using a total of 11 blocks of H&E-stained images and 10 blocks of IHC-stained images.

### 3.1. Registration

The staining method had little effect on the quality of the automatic registration (Figure 3). Registration of the H&E images decreased MSE by 57.14 ± 65.61 (p << 0.001, n = 121) and increased the SSIM by 0.03 ± 0.03 (p << 0.001, n = 121) (Figure 3, top). For the IHC images, registration decreased MSE by 58.86 ± 43.10 (p << 0.001, n = 112) and increased SSIM by 0.09 ± 0.04 (p << 0.001, n = 112) (Figure 3, bottom).

**Fig. 3:**
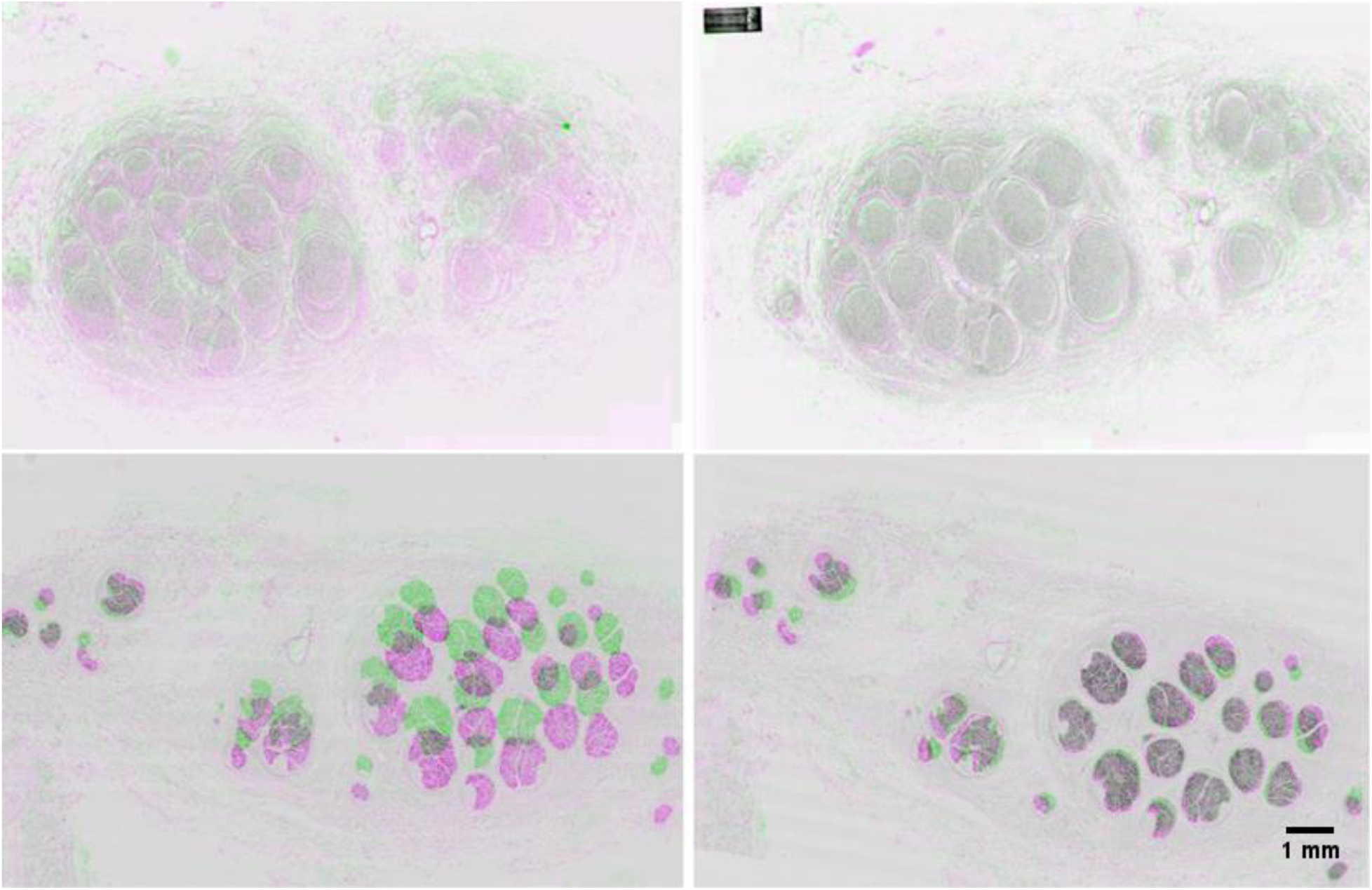
A comparison of H&E images (top) and IHC images (bottom) before (left) and after (right) automatic registration. Areas of pink and green show regions of dissimilarity, whereas grey shows aligned regions.

### 3.2. Fascicle Detection

The detection step for H&E slices produced a set of fascicle bounding boxes for each image (Figure 4). The network trained on the original images had a precision, recall and F1-score of 0.92, 0.90, and 0.91, respectively. The network trained on the registered images performed comparably, with precision, recall and F1-score of 0.91, 0.92 and 0.91, respectively. Both networks were tested in the subsequent segmentation step. In contrast, the circle detection method performed worse, with precision, recall and F1-scores of 0.19, 0.45, and 0.27, respectively. A separate detection step was not required for IHC slices.

**Fig. 4:**
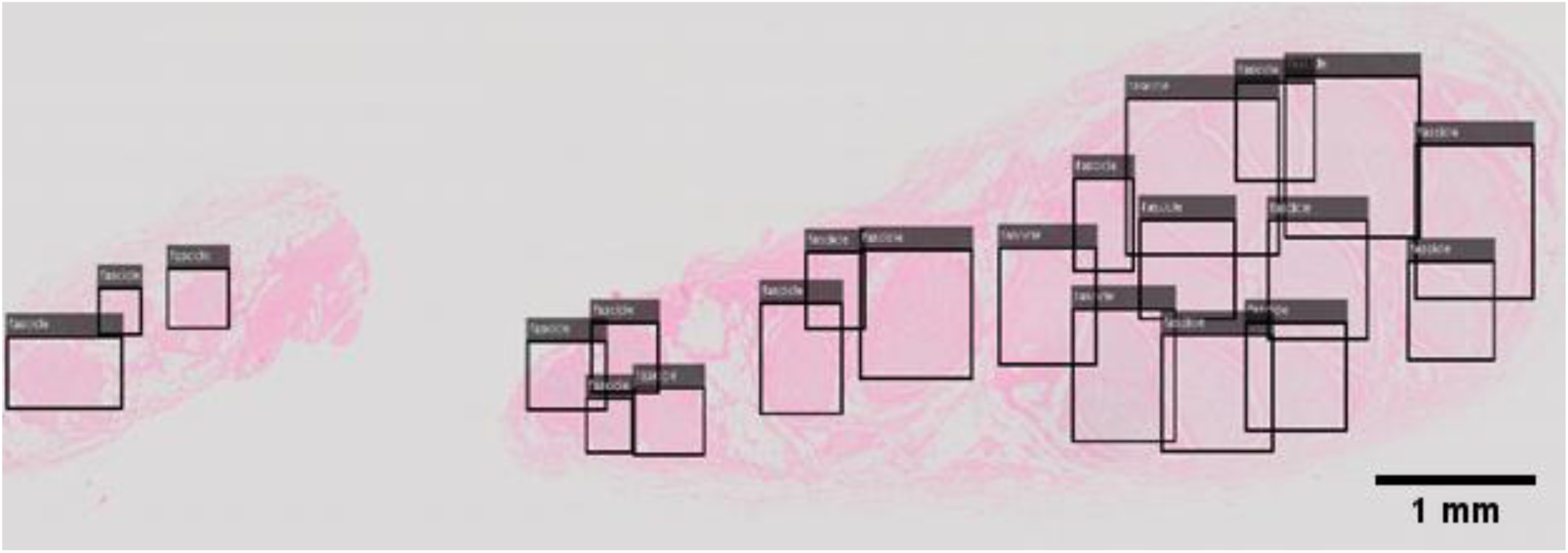
Bounding boxes indicating the fascicle sizes and locations automatically detected using the RCNN, on a sample histological slice.

### 3.3. Fascicle Segmentation

The segmentation outputs varied according to the method implemented (Figure 5, Table 2). Active contour segmentations initialized using the neural network trained on the original, unregistered images showed the highest performance on all three segments, as well as overall, with an IOU of 0.61±0.03. These segmentations were used for reconstruction. Otsu’s method with follow-up morphological processing segmented the IHC slices well, with an IOU of 0.900±0.093.

**Fig. 5:**
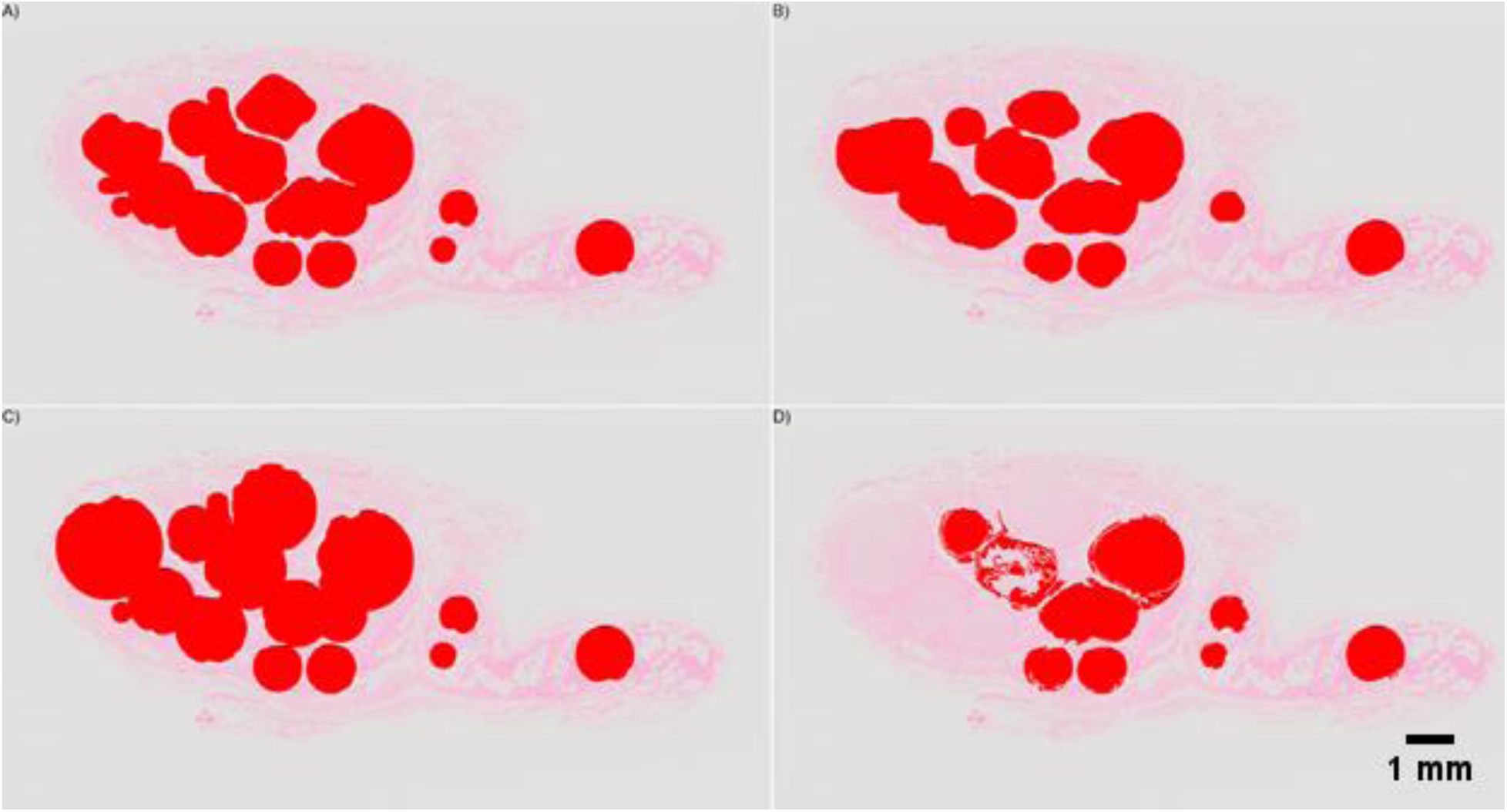
Images illustrating the segmentation of individual fascicles from histological slices. Segmentations (red) are overlaid on the original H&E image. A) Outside-in active contours B) Inside-out active contours C) Otsu’s Method D) K-means clustering.

**TABLE 2.**
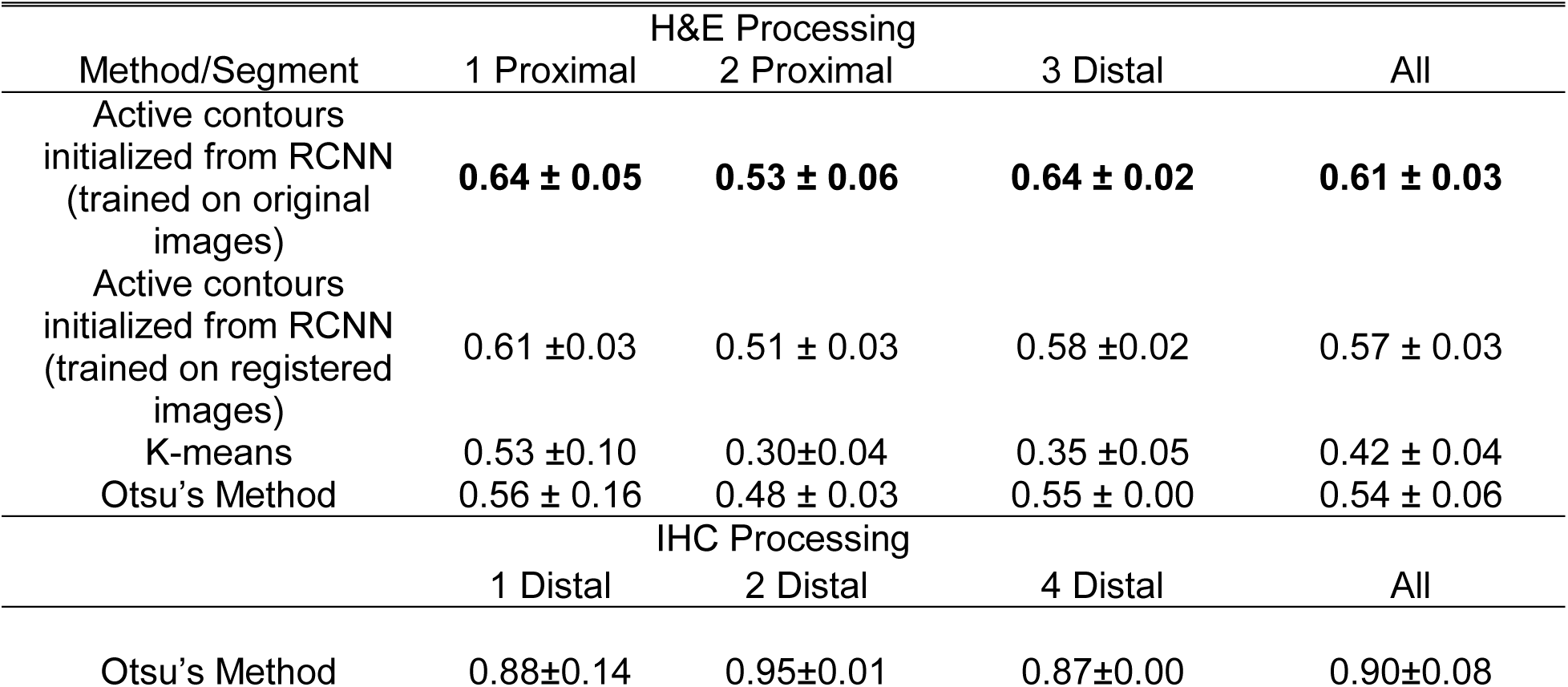
Mean IOU±SD determined for each of the segmentation methods implemented on images from each staining method.

| Method/Segment | H&E Processing |  |  |  |
| --- | --- | --- | --- | --- |
|  | 1 Proximal | 2 Proximal | 3 Distal | All |
| Active contours initialized from RCNN (trained on original images) | **0.64 ± 0.05** | **0.53 ± 0.06** | **0.64 ± 0.02** | **0.61 ± 0.03** |
| Active contours initialized from RCNN (trained on registered images) | 0.61 ± 0.03 | 0.51 ± 0.03 | 0.58 ± 0.02 | 0.57 ± 0.03 |
| K-means | 0.53 ± 0.10 | 0.30 ± 0.04 | 0.35 ± 0.05 | 0.42 ± 0.04 |
| Otsu's Method | 0.56 ± 0.16 | 0.48 ± 0.03 | 0.55 ± 0.00 | 0.54 ± 0.06 |
|  | IHC Processing |  |  |  |
|  | 1 Distal | 2 Distal | 4 Distal | All |
| Otsu's Method | 0.88 ± 0.14 | 0.95 ± 0.01 | 0.87 ± 0.00 | 0.90 ± 0.08 |

### 3.4. Reconstruction

The fully automatic, semi-automatic, and extruded models had similar IOUs, but qualitatively differed in appearance (Figures 6 and 7, Table 3). EX had higher IOUs than the fully automatic models for ten of the twelve examined blocks. Of the semi-automatic models, MRAS models performed superior to EX and ARMS models. For H&E images, reconstruction with hole-fixing of discontinuous fascicles marginally improved IOUs, while the opposite was true for IHC images. In most models, the quantitative effect of hole-fixing was minor; the greatest impact occurred in the third distal block of the second specimen, with a 1.88% increase in IOU (Figure 8).

**Fig. 6:**
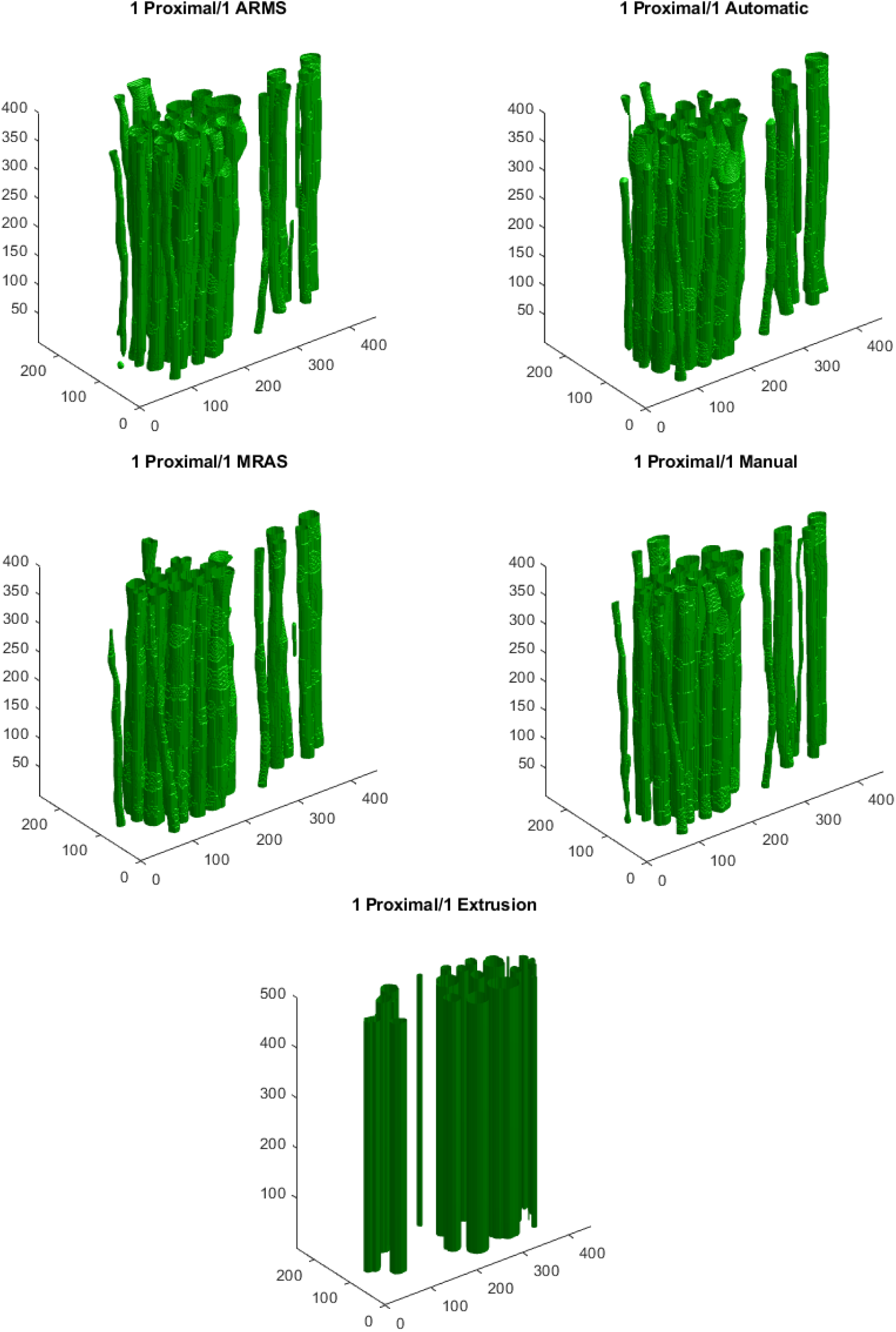
Models generated from the first proximal block of the first nerve specimen, stained with H&E. Shown are the fully automatic, semi-automatic, and fully manual models, along with an extrusion using the first slice of the block. Dimensions on each axis are expressed in pixels.

**Fig. 7:**
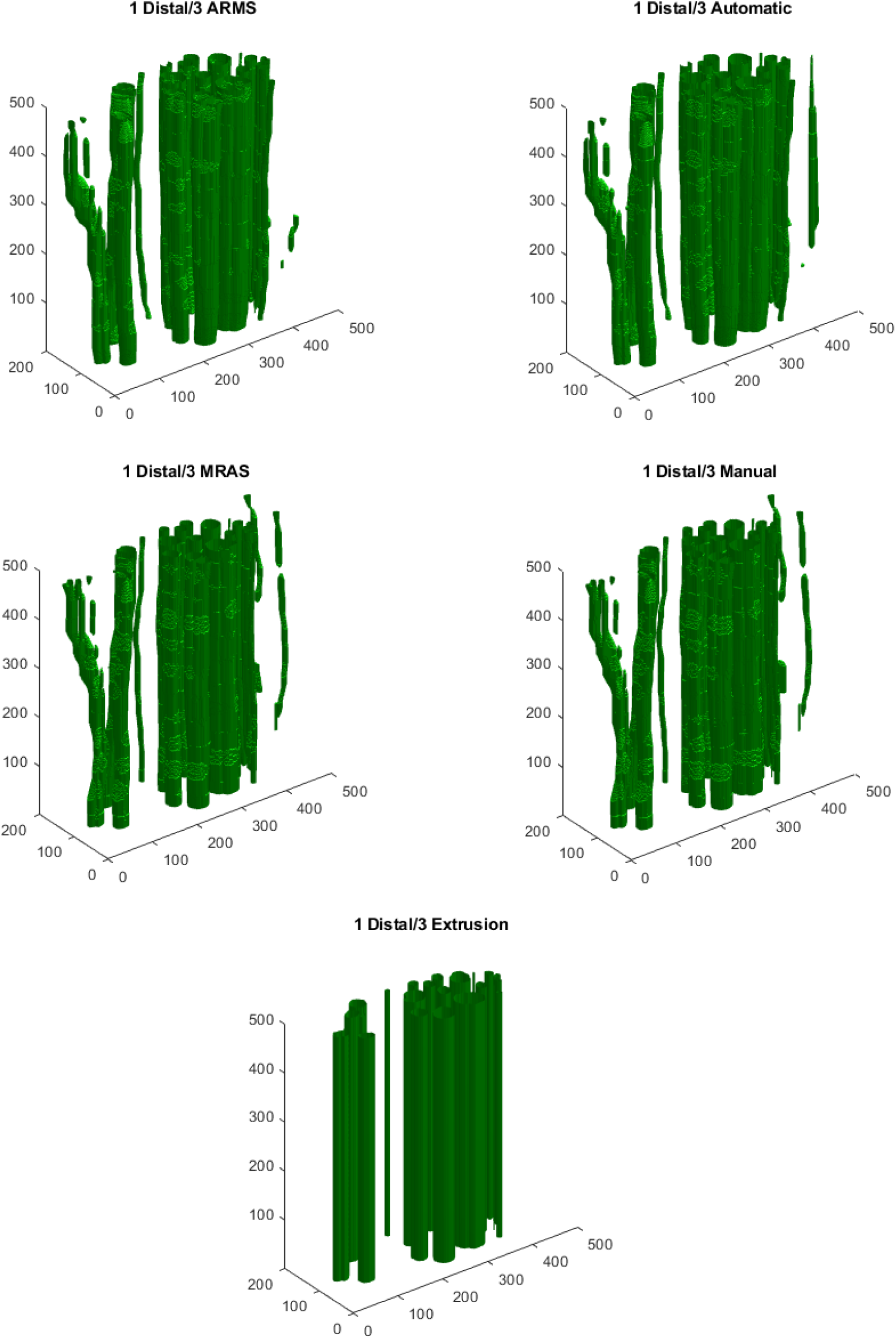
Models generated from the third distal block of the first nerve specimen, stained with IHC. Shown are the fully automatic, semi-automatic, and fully manual models, along with an extrusion using first slice of the block. Dimensions on each axis are expressed in pixels.

**Fig. 8:**
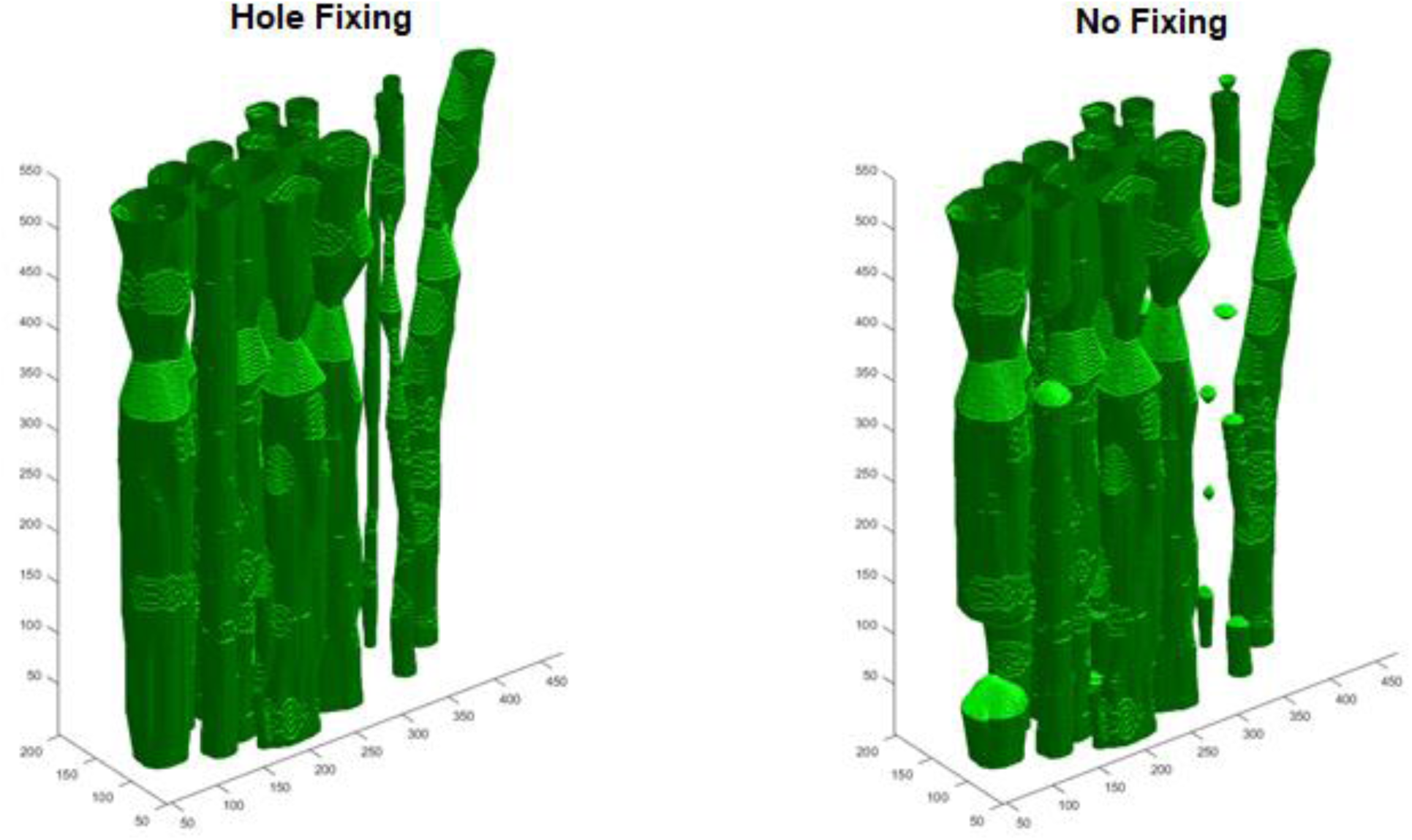
The effect of hole-fixing. Note that the connected fascicles in the fixed image (left) are separate in the unfixed image (right). Dimensions on each axis are expressed in pixels.

**TABLE 3.** Mean IOU±SD of the different reconstruction methods for H&E and IHC images, compared to fully manual reconstructions. The two semi-automatic methods are listed separately as they were not directly compared to the other three methods, and instead were used to identify contributions of registration and segmentation to the final IOU scores.

| Method | Mean IOU<br>(H&E)<br>n=6 | Mean IOU<br>(IHC)<br>n=6 |
| --- | --- | --- |
| HF | 0.426±0.062 | 0.398±0.149 |
| NF | 0.423±0.064 | 0.399±0.150 |
| EX | 0.485±0.110 | 0.528±0.064 |
| ARMS | 0.476±0.108 | 0.400±0.150 |
| MRAS | 0.612±0.066 | 0.900±0.093 |

## 4. DISCUSSION

The goal of this project was to create an algorithm that can automatically reconstruct fascicular branching patterns from consecutive cross-sections of peripheral nerves. The current pipeline intends to alleviate the need for time intensive manual processes and facilitate the construction of computational models for neuroprosthetic and neuromodulation applications. The pipeline successfully generated models from images based on both H&E- and IHC-stained histological slices. Contrary to expectation, some models produced using our fully automatic method were quantitatively poorer than those based on extrusion. However, fully automatic models provided qualitative information about fascicle shape and branching patterns that were lost using extrusions. The generation of semi-automatic models, ARMS and MRAS, demonstrated the impact of the registration and segmentation on the final reconstruction.

### 4.1 Registration

Registration improved the overall alignment of sequential slices. The improved alignment was demonstrated by the average decrease and increase in MSE and SSIM, respectively. However, unexpectedly high standard deviations for both MSE and SSIM indicated that not all pairs of consecutive images benefitted equally from registration. In some cases, registration quantitatively decreased the alignment, likely due to the emphasis on aligning the fascicles and decreased visibility of the connective tissue and nerve boundary caused by preprocessing. Although the study that introduced this four-step registration method mentioned no such difficulties (22), the current study indicates the need to more closely evaluate the registration outcomes at each step. Finally, the registration method tended to align epineurium boundaries, thus potentially straightening out the epineurium of successive slices relative to actual *in vivo* geometry. This was not a key concern for the neuroprosthetic context, since implantation of certain neural interfaces such as nerve cuff electrodes may straighten out the nerve regardless. However, more generally, a straightened nerve may not represent the nerve course *in situ*. An external reference, generated using MRI or MicroCT/OPT, could be used to acquire pre-histological external boundary information, to which the algorithm could be modified to conform.

### 4.2 Fascicle Detection

The RCNN successfully detected fascicles in the H&E slices, as demonstrated by the high F1-score. Investigating neural network architectures other than VGG-16 (e.g., YOLOv3 (28)) may further improve detection performance; however, fascicle detection was likely not the bottleneck in our current pipeline, as registration and segmentation proved more challenging. While unsupervised machine learning methods have been used previously for fascicle detection, our study is the first to use a trained RCNN to detect fascicles in H&E stained images (29). Additionally, we specifically and uniquely quantified the performance of fascicle detection to provide a point of comparison for future implementations.

### 4.3 Fascicle Segmentation

The accuracy of fascicle segmentation depended on staining method, and thus the amount of pre- and post-processing involved with the images. Our fully automatic method achieved accuracies ranging from 53 to 64% for the H&E images. Although our segmentation scores for H&E images were lower than the 89 to 94% reported for semi-automatic methods (29), our fully automatic method is more easily scalable for generating models from multiple individuals. In contrast, our segmentation of IHC images achieved 90% accuracy using Otsu’s method with minimal pre and post-processing. The performance on IHC images was comparable to the 94.5% similarity reported for active contour segmentation of Micro-CT slices, relative to manually labeled boundaries (16). Active contour-based segmentation provided good approximation of fascicle boundaries, but could miss small or less visible fascicles.

Some difficulty was experienced when segmenting densely packed fascicles, which we partially addressed using watershed splitting. A potential alternative to active contours would be to train a semantic segmentation deep neural network. Semantic segmentation involves using a CNN in a pixel-wise manner, in our case classifying each pixel as part of the fascicle or the background. If supplied with sufficient training data, this could eliminate the detection step and improve the segmentation performance. Although a more expensive preparation, IHC stained nerves did not require advanced image processing. Regardless of the staining method used, the segmentation quality of individual slices markedly affected reconstruction outcomes.

### 4.4 Reconstruction

Several mechanisms contributed to discontinuous fascicles due to the logic of the reconstruction method: 1) the registration could not align the fascicle; 2) the fascicle was not detected in a particular slice(s); 3) the fascicle was not fully segmented (i.e., no overlap with neighbouring slice); or 4) the fascicle experienced an abrupt change in position. Hole-fixing appeared to help in some cases, but hindered reconstruction in others. For H&E slices, hole-fixing tended to introduce a quantitatively minor benefit in terms of IOU. The effect was opposite for IHC slices, where blocks without hole-fixing showed a minor increase in IOU. This could be due to the high quality of segmentation for IHC slices. With few missing fascicles requiring compensation, applying additional hole-fixing would likely introduce more errors. Another possibility is that our algorithm generally performs well without hole-fixing when dealing with large, well-defined fascicles. Thus, hole-fixing mostly benefits small fascicles and only minor changes in IOU would occur after its application. Whether or not to use hole-fixing may warrant a case-by-case examination, especially since reconstruction is one of the least time-intensive components of the overall pipeline.

### 4.5 Impact of Registration and Segmentation on Reconstruction Quality

Using our semi-automatic models, we could ensure a good registration or segmentation and thus better understand how each of these two steps contributed to the IOU scores after reconstruction. In some cases, IOU demonstrated that simple extrusions were superior to our fully automatic method, but only in one case was the extrusion superior to MRAS semi-automatic slices. This suggests the importance of registration in determining the final reconstruction IOU. Additionally, the possibility exists that the fascicles within the automatic models appear collectively displaced (within the global reference frame) relative to those of the manual models. This lack of alignment in the global reference frame may cause low IOU scores, although the fascicles are correctly aligned relative to one another. Therefore, compared to segmentation, registration carries larger impact on the final quality of reconstruction.

### 4.6 Outlook

Reviewing all blocks where extrusions were superior to the automatic method identified potential causes for variation in IOU scores. Specifically, we noted the following possible causes: local misalignment in registration (8 cases); large fascicles missing (i.e., detection issue; 2 cases); small fascicles missing (i.e., segmentation issue; 5 cases); discontinuous fascicles in manual segmentation (3 cases); and high fascicle density (1 case). For registration, increasing the contrast of fascicles in pre-processing or increasing the focus on external nerve boundaries, could create closer-to-manual models. Adding an extra registration at low resolution could increase the focus on alignment of exterior boundaries. In three cases, fascicles rapidly changed position and appeared discontinuous in the fully manual nerve slices. Taking one slice every 100µm would allow for more precise tracking of individual fascicles along the length of the nerve, reducing the likelihood of discontinuities caused by rapid changes in fascicle position. Detection could be improved by acquiring more nerve slices to increase the amount of training data. Adding a minimum size constraint could improve segmentation, ensuring that fascicles detected in the mask are not lost after outside-in segmentation.

### 4.7 Contributions to Literature

The current work presents a unique fully automatic 3D reconstruction method. We quantified the performance of each step of our process in order to establish a standard for future explorations of this technology. Furthermore, except for one group that created their own visualization software (18), others used commercial reconstruction software to generate their models, reducing the accessibility to their methods(16,17,19,29). Every step of our method was implemented in MATLAB, which both simplified the processing pipeline and reduced the barrier to entry. While further work remains to achieve a completely reliable process, the modular nature of our method will make it easy to improve individual steps in the pipeline. Moreover, modularity makes it possible to independently implement any step within semi-automatic workflows.

## 5. CONCLUSION

Currently existing peripheral nerve computational models for neural interfaces predominantly use simplified neural anatomy. Previous research has shown that the conclusions drawn from computational models can differ depending on the level of anatomical detail in the model; however, the construction of anatomically accurate models is very time consuming when done manually. We introduced a framework to automatically generate nerve models based on serial histological cross-sections. While models could be produced from both H&E and IHC slices, the easier processing and superior quality resulting from the IHC slices suggests that this avenue should be preferred. While improvements are still required, this study provides a baseline and stable platform for future development of algorithms to generate accurate computational models to support the development of neural interfaces.

## Abbreviations

ARMS: Automatic Registration, Manual Segmentation
MRAS: Manual Registration, Automatic Segmentation

## Conflicts of Interest

The authors have no conflicts of interest to report.

## ACKNOWLEDGEMENTS

This project was supported by the National Sciences and Engineering Research Council of Canada (RGPIN-2014-05498) as well as the Institute of Biomaterials and Biomedical Engineering at the University of Toronto. The authors would also like to acknowledge Milan Ganguly of the Centre for Phenogenomics (Toronto, Canada) and his team for their help in providing the data used in this project.

